# A system-level view on the function of natural eukaryotic biomes through taxonomically resolved metabolic pathway profiling

**DOI:** 10.1101/2022.07.27.501711

**Authors:** Jayson Gutierrez, Pascal I. Hablützel

**Author notes:** **Corresponding author:** Jayson Gutierrez. **Competing Interests:** The authors declare no competing interests.

## Abstract

High-throughput sequencing of environmental samples has dramatically improved our understanding of the molecular activities of complex microbial communities in their natural environments. For instance, by enabling taxonomic profiling and differential gene expression analysis, microbiome studies have revealed intriguing associations between community structure and ecosystem functions. However, the effectiveness of sequence data analysis to characterize the functioning of microbial ecosystems at the systems level (*e*.*g*. metabolic pathways) and at high taxonomic resolution has thus far been limited by the quality and scope of reference sequence databases. In this work, we applied state of the art bioinformatics tools to leverage publicly available genome/gene sequences for a wide array of (mostly eukaryotic) planktonic organisms to build a customized protein sequence database. Based on this, our goal is to conduct a systems-level interrogation of environmental samples, which can effectively augment the insights obtained through traditional gene-centric analysis (*i*.*e*. analysis of single gene expression profiles at the genome-wide level). To achieve this, we utilized the popular HUMAnN pipeline, which has proven effective at delineating taxon-specific metabolic pathways that may be actively contributing to the overall functioning of a microbiome. To test the efficacy of our database customization for mapping metabolic pathway activities in complex planktonic ecosystems, we reanalyzed previously published metatranscriptome datasets derived from different marine environments. Our results demonstrate that database customization can substantially improve our ability to quantitatively assess core metabolic processes across taxonomically diverse marine microbiomes, which have so far remained largely uncharacterized at the systems level. By further expanding on the taxonomic and functional complexity of our database with newly released high-quality genome assemblies and gene catalogs for marine microbes, we aim to improve our ability to map the molecular traits that drive changes in the composition and functioning of marine planktonic networks through space and time.

## INTRODUCTION

Natural microbial communities fulfill many essential ecosystem functions [1]. They contribute about half of global primary productivity through photosynthesis [2], drive global biogeochemical cycles [3, 4], catalyze bioremediation [5], and safeguard ecosystem stability through multifarious species interactions [6]. Such communities are often species-rich and dynamic through time [7]. These characteristics raise fundamental questions in ecological and evolutionary research: How do species-rich natural microbial communities organize themselves, how do they achieve dynamic stability and how much is ecosystem function related to species diversity? But dynamic equilibria of complex communities are difficult to reproduce under controlled laboratory conditions and many species still cannot even be cultured in the lab. When controlled experimental approaches fail, field observations are valuable means to provide critical insights into the ecological dynamics of microbial ecosystems [8]. To this end, the application of next-generation sequencing (NGS) technologies has fueled the use of global gene expression profiles from environmental samples as an approximation of ecological function in microbial communities [9]. The approach has become a standard procedure to interrogate the physiological status of a variety of microbiomes from disparate ecosystems, such as terrestrial [10, 11], aquatic [12–14] and human gut environments [15, 16].

Metabolic activity of a microbial community can be investigated compositionally by modeling gene-expression. Multi-species gene-expression networks can subsequently be decomposed to reconstruct metabolic pathways of single species. This second step is highly dependent on the taxonomic resolution of the reference database [17–20]. Therefore, the success of this type of modeling depends on the availability of high quality annotations for full genome assemblies or nearly complete metagenome assembled genomes (MAGs), which limits its applicability to microbiomes with well characterized organisms. One way to partially overcome this limitation is by exploiting large databases of functionally annotated sequence data of related species. This information source allows us to estimate well known molecular pathways that sustain core cellular processes (*i*.*e*. biosynthesis and catabolism). A popular software tool that enables a systems-level view of microbiomes is HUMANnN [21, 22], which provides a convenient, automated way for mapping sequence reads to a well curated database of metabolic pathways, *e*.*g*. the MetaCyc database [23]. Microbiome analysis software tools such as HUMANnN have proven quite effective at mapping short sequence reads to taxon-specific gene families when applied on metagenome and metatranscriptome datasets derived from comprehensively studied environments using the default databases (ChocoPhlAn pangenome and UniRef50/90 databases) [22, 24]. Yet, it remains to be seen the performance of those tools when applied on samples taken from largely unexplored environments, where the microbiomes of which are likely to exhibit unique patterns of molecular sequence diversity poorly represented in existing reference sequence databases [24–27]. Recent studies have raised awareness on the necessity of database customization for improving our ability to characterize the metabolism of yet underexplored microbiomes [24, 28].

Marine planktonic ecosystems encompass one of the most taxonomically and ecologically diverse microbiomes, which in addition to sustaining the whole marine food web also play critical roles in global biogeochemical cycling, including carbon sequestration in deep-sea sediments [29–32]. Most meta-omics studies of the marine microbiome have so far placed emphasis on the reconstruction of extensive gene catalogs and the delineation of protein functional clusters from a large set of samples collected across different oceanic provinces [33–35]. Notably, such sequence catalogs have proven effective as a general framework for studying planktonic ecosystems, including the association between gene sequence diversity and expression patterns with ecoregions, metabolic processes and trophic modes [13, 14, 25, 26]. Although these bioinformatic approaches have greatly advanced our understanding of planktonic ecosystems, novel bioinformatics tools facilitating a systems biology analysis of meta-omics datasets are needed for *e*.*g*. adequate parameterization of integrative ecosystem biology models, which could offer detailed mechanistic explanations of trophic interactions and biogeochemical cycling [36–41].

In this work, we aim to demonstrate that custom sequence databases enable detailed functional profiling of (eukaryotic) plankton communities. Specifically, we aimed at obtaining fine-grained taxonomically resolved activity of metabolic pathways. To do so, we leveraged publicly available whole-genome assemblies and catalogs of predicted gene sequences for a wide array of planktonic organisms. Subsequently, we deployed the HUMANnN pipeline (3.0) [21, 22] to quantify metabolic pathway abundance (a proxy for activity). Our motivation for exclusively building a HUMANnN-compatible database was twofold: 1) this tool is quite flexible, offering the users the possibility to efficiently explore large meta-omics datasets from underexplored environments under a great variety of settings, including different identifiers for protein and metabolic pathway annotations; and 2) this tool enables the automated computation of rough estimates about gene expression patterns aggregated at the metabolic pathway level, which can facilitate a systems level interrogation of the physiological activity of microbial communities. This latter point is, arguably, one of the most critical aspects of a systems biology-oriented bioinformatics pipeline. To assess the efficacy of our database customization we reanalyzed previously published metatranscriptome datasets derived from samples taken across different marine environments.

## MATERIALS & METHODS

### Data provenance and taxonomic coverage

To build the plankton-specific database, we initially gathered relevant taxonomic information by querying The World Register of Marine Species [42]. Importantly, we narrowed down our search for plankton organisms reported in and around the Belgian part of the North Sea (our main study area). Based on an initial set of 1103 entries retrieved from the search, we then queried the NCBI repository for relatively high quality genome assemblies (minimal N50 = 1000) available for plankton organisms with taxonomic description at the species level. This resulted in a list of 365 genomes for Eukaryota (249), Bacteria (115) and Archaea (1). To further expand the functional diversity and taxonomic representation of our database, we re-annotated sequences derived from large publicly available catalogs reported for marine plankton species, including well curated single-cell and metagenome assembled genomes (SMAGs and TOPAZ) collected during the Tara Oceans expedition [43, 44]. Further, we also extracted sequences from the EukZoo database [45], which was obtained by annotating sequences from the Marine Microbial Eukaryotes Transcriptome Sequencing Project (MMETSP) and various other genomes and transcriptomes of organisms that were not included in the MMETSP [46]. Importantly, for these catalogs we focused on sequences that had at least taxonomic affiliations at the family level.

### Functional annotation

Prokaryotic genomes were processed via the Struo2 pipeline [24], which relies on prodigal as a gene calling tool [47], and the DIAMOND (version 0.9.18) alignment tool [48] to annotate sequences against a recent release of the UniRef90 database [49] that encompasses >130M protein families (version 2020-01) in the form of clustered sequences based on similarity at 90%. On the other hand, the set of eukaryotic genomes obtained through the search across NCBI were reannotated at the structural and functional level. For structural annotation (*ab initio* gene prediction) of eukaryotic genomes we utilized the software BUSCO [50] to obtain training sets in the form of candidate stretches of DNA sequences in contigs/scaffolds that might encode for a functional protein. These BUSCO-generated hints were then used for more refined gene prediction tasks using Augustus [51], which yielded a fasta file with a list of candidate genes. Similar to the functional annotation of the prokaryotic gene set described previously, the total set of eukaryotic genes collected were annotated based on sequence similarity using DIAMOND [48] and the UniRef90 database [49] as a reference (see supplementary information for details). The resulting set of sequences for both prokaryotic and eukaryotic organisms pooled together from these different sources (3803430 sequences) was then clustered into protein families using the MMseqs2 command line tool (at 90% sequence identity, with minimal fraction of aligned residues set at 0.8) [52]. This yielded a less redundant and slightly more compact representation of our database comprising a total of 2.867.766 protein families variably allocated across most major taxonomic groups of eukaryotic plankton organisms (Figure 1), in which ∼94% (2682205) of the sequences in our database are allocated.

**Figure 1.**
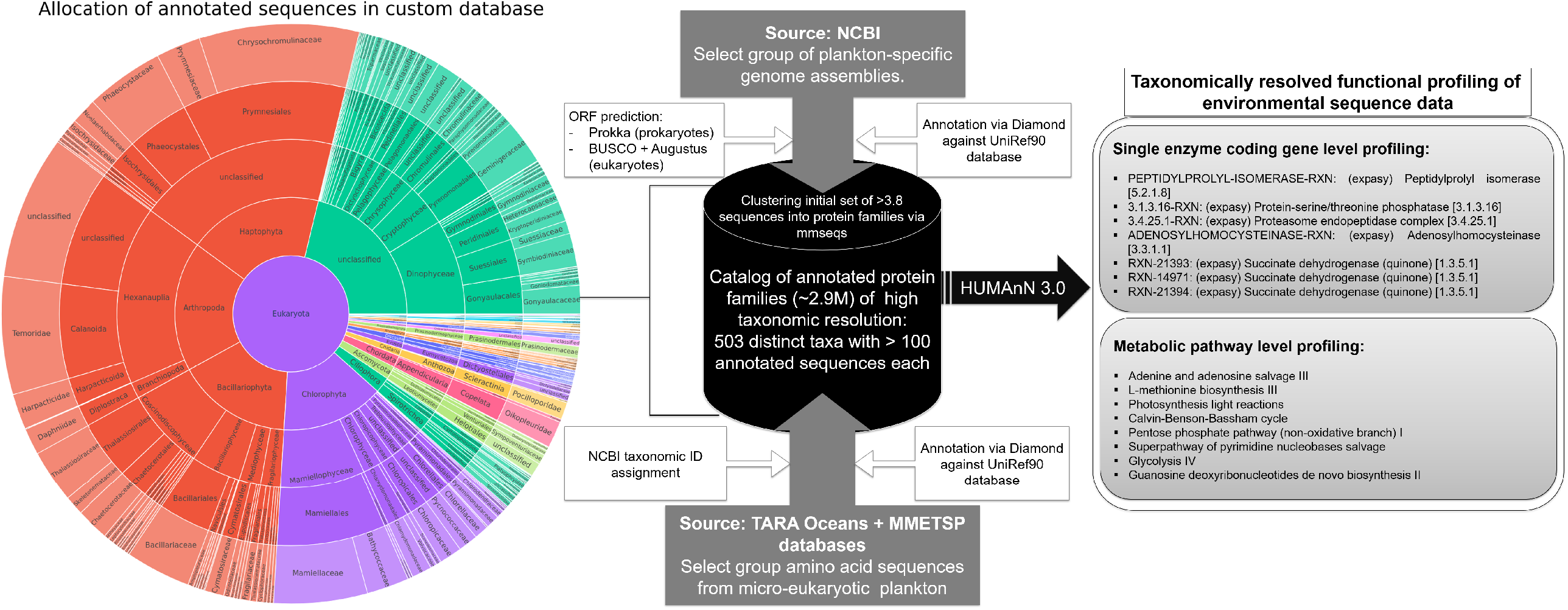
Summary of our workflow devised to create the plankton-specific database of annotated protein families. An initial list of >6M candidate gene sequences was created by mining and processing sequence information from various publicly available sequence resources (see Materials and Methods). This involved fetching plankton-related predicted protein sequences, as well as genome assemblies that were subjected to ORF prediction using popular bioinformatics tools (i.e. Prokka, BUSCO and Augustus). UniRef90 annotations were then transferred to sequences found as relatively high-quality alignments using Diamond (see Materials and Methods). This derived in >3.8M annotated protein sequences, which were then clustered into protein families, using the mmseqs tool, to eliminate redundancy and keep our database light with the aim of optimizing for query times when running the profiling tool HUMAnN. Our custom database encompasses a wide variety of distinct planktonic taxonomic groups (sunburst plot on the left), which enable us to profile at high taxonomic resolution the activity of plankton communities in terms of single enzymes and metabolic pathways (panels on the right) using HUMAnN.

### Quality control of Metatranscriptomics data

Each metatranscriptome analyzed here was subjected to quality control using the BBTools suit, a set of fast, multithreaded bioinformatics tools designed for the analysis of DNA and RNA sequence data [53]. Specifically, we use the bbduk tool to conduct adapter trimming (with parameters: ktrim=r k=27 hdist=1 tpe tbo -eoom), quality control trimming (with parameters: qtrim=r trimq=20 -eoom), quality filtering to discard reads with average quality below 20 (with parameter: maq=20). Moreover, sequence reads with length < 50 nucleotides were discarded using a custom python script. Finally, custom python scripts relying on the Biopython package [54] were used to conduct all the necessary bioinformatics tasks described above.

### Case study 1: Elferink et al. (2020)

To assess the potential of our database for meta-omics profiling of plankton communities we reanalyzed previously published metatranscriptome datasets derived from different marine regions. Firstly, we examined 12 samples (6 samples for each area, two sampling sites for area, 3 replicates per site) taken from coastal surface waters of Greenland and Iceland [55] by running HUMANnN with default settings, using as reference the natively installed databases (ChocoPhlAn/UniRef90), which cover a wide variety of distinct taxonomic groups [21, 22]. It should be noted that we prioritized functional profiling of environmental samples against reference sequences with a minor degree of uncertainty regarding their potentially encoded function, as specified by the UniRef90 database. However, profiling results are likely to improve substantially by applying HUMANnN using the UniRef50 database as a reference, which clusters sequences into more broadly defined protein families with larger sequence dissimilarity [49]. The same logic applies for the remaining case studies described below.

### Case study 2: Berg et al. (2018)

To further demonstrate the potential of our database customization to improve our ability to characterize the physiological status of plankton communities using gene expression patterns as a proxy, we examined additional publicly available metatranscriptome datasets. For instance, we profiled numerous samples taken in the Baltic Sea during a cyanobacterial bloom [56].

### Case study 3: Kolody et al. (2019)

We also re-examined a time series metatranscriptome dataset collected from North pacific high nutrient, low-chlorophyll waters, which were previously analyzed for diel expression patterns in response to nutrient gradients and viral infections [57].

## RESULTS

### Case study 1: Elferink et al. (2020)

Although in a previous study HUMANnN has proven effective at mapping major metabolic activities in samples taken from a different marine environment [21], we found that for this particular set of samples and under default settings, HUMANnN performs poorly at mapping sequence reads to enzyme-coding sequences (labeled with EC numbers) involved in several metabolic functions (Figure 2A). In stark contrast, we found that HUMANnN can be more than 2 orders of magnitude more effective at mapping reads to EC numbers when running it through our customized, plankton-specific database (Figure 2A). Furthermore, when examining the activity of entire metabolic pathways, which cover a variable number of functionally related enzymes, we observed contrasting patterns between Arctic (Greenland stations) versus Subarctic coastal waters (Iceland stations), and even between sampling sites within the same region (Figure 2B). For instance, we found that gene expression levels associated with core metabolic processes (glycolysis, pentose phosphate pathway, and light and dark reactions) biosynthesis of nucleotides, amino acids, lipids, and carbohydrates are relatively highly expressed, which show detectable activity patterns across numerous species. However, variation can exist between sampling sites within the same regions, especially within the Greenland coast. Although sequencing errors may explain part of this variation, environmental gradients within the same region cannot be ruled out as important drivers of such differences in metabolic pathway activity. More generally, gene expression differences across less active metabolic pathways involved in a variety of catabolic and anabolic processes seem to exist between plankton communities observed in Greenland and Island stations, which are likely to be less sensitive to differences in environmental variables. Overall, these results indicate that active metabolic pathways are widely distributed across numerous species, which tend to variably contribute to the bulk of essential biosynthetic activities in the community. These findings are in line with the original study. On the other hand, analysis of the dominant planktonic groups in these surface waters, namely dinoflagellates (Dinophyta) and diatoms (Bacillariophyta), suggests that these organisms tend to exhibit site-specific activity for pathways involved in primary metabolism (Figure 2C), which provides a broader view on the global metabolic activity described in the original study [55]. Overall, these results suggest the presence of metabolic niches for broadly defined taxonomic groups, which are likely set by differences across sampling sites in terms of temperature, salinity and nutrient availability.

**Figure 2.**
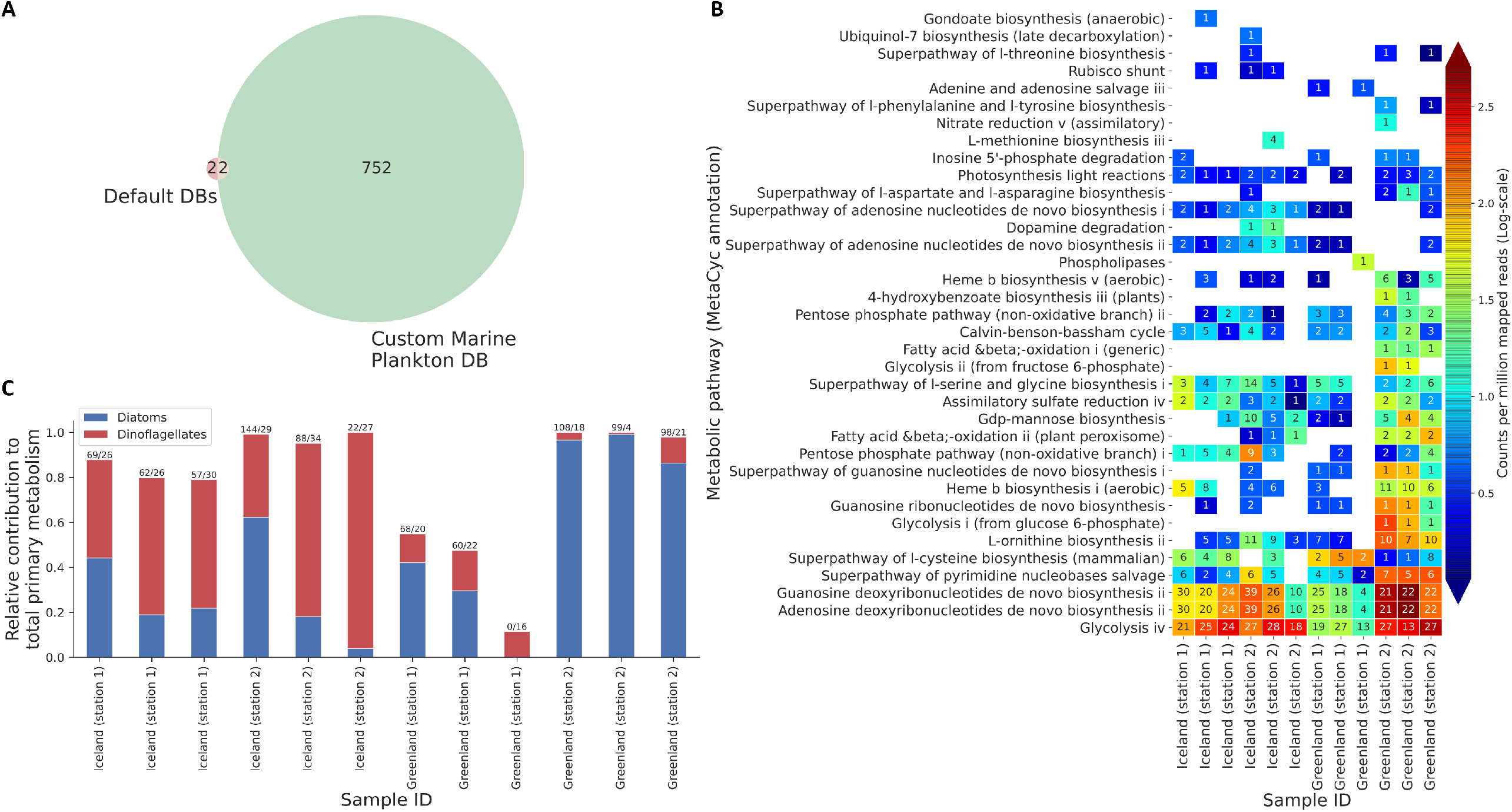
A, Venn diagram summarizing the total number of enzymes (identified by their E.C.) that RNA sequence reads in the samples analyzed (BioProject record PRJEB37134) were mapped through HUMAnN using default databases (DBs) versus our plankton-specific database. This diagram illustrates results taking together all the samples analyzed. B, heatmap illustrating aggregated abundance across contributing species (indicated in each cell of the grid) to a specific metabolic pathway, for the distinct geographical locations (stations) the samples were taken from. Importantly, metabolic pathway abundance scores (a proxy for activity in the context of metatranscriptome data) were normalized (cross-sample) as counts per million mapped reads. C, bar chart summarizing the relative contribution of diatoms versus dinoflagellates to primary metabolic activity in the plankton communities analyzed. Numbers shown on top of each bar give the ratio of the total number of diatom to dinoflagellate species found to contribute to primary metabolism.

### Case study 2: Berg et al. (2018)

Overall, our results confirm the dominant metabolic activity of Cyanobacteria (Figure 3A), and also reveal several other major planktonic groups, including Proteobacteria, Chlorophyta, Dinophyceae and Ciliophora, which seem to exhibit variable metabolic activity via specific pathways across the different samples examined (Figure 3A). In congruence with previous observations, we found that Cyanobacteria dominated metabolic activity through photosynthesis light reactions, CO_2_ fixation and energy production pathways, and surprisingly enough via the Rubisco shunt pathway, which shows a high efficiency of carbohydrates-to-oil conversion (Figure 3B). Interestingly, we found that within a given sampling station (Stockholm archipelago) global metabolic pathway activity profiles across the plankton community tend to show quite similar patterns throughout the different seasons examined [56] as revealed by the clustergram (Figure 3B). Moreover, the other planktonic groups found across the different samples analyzed predominantly showed relatively high activity in ATP synthesis (Figure 3B). Moreover, within the Cyanobacteria group we found a great variety of species contributing to the different metabolic activities described above, with Cyanobium being the dominant metabolically active species, with other species and strains mostly associated with the Synechococcus genus variably contributing different metabolic functions (Figure 3C).

**Figure 3.**
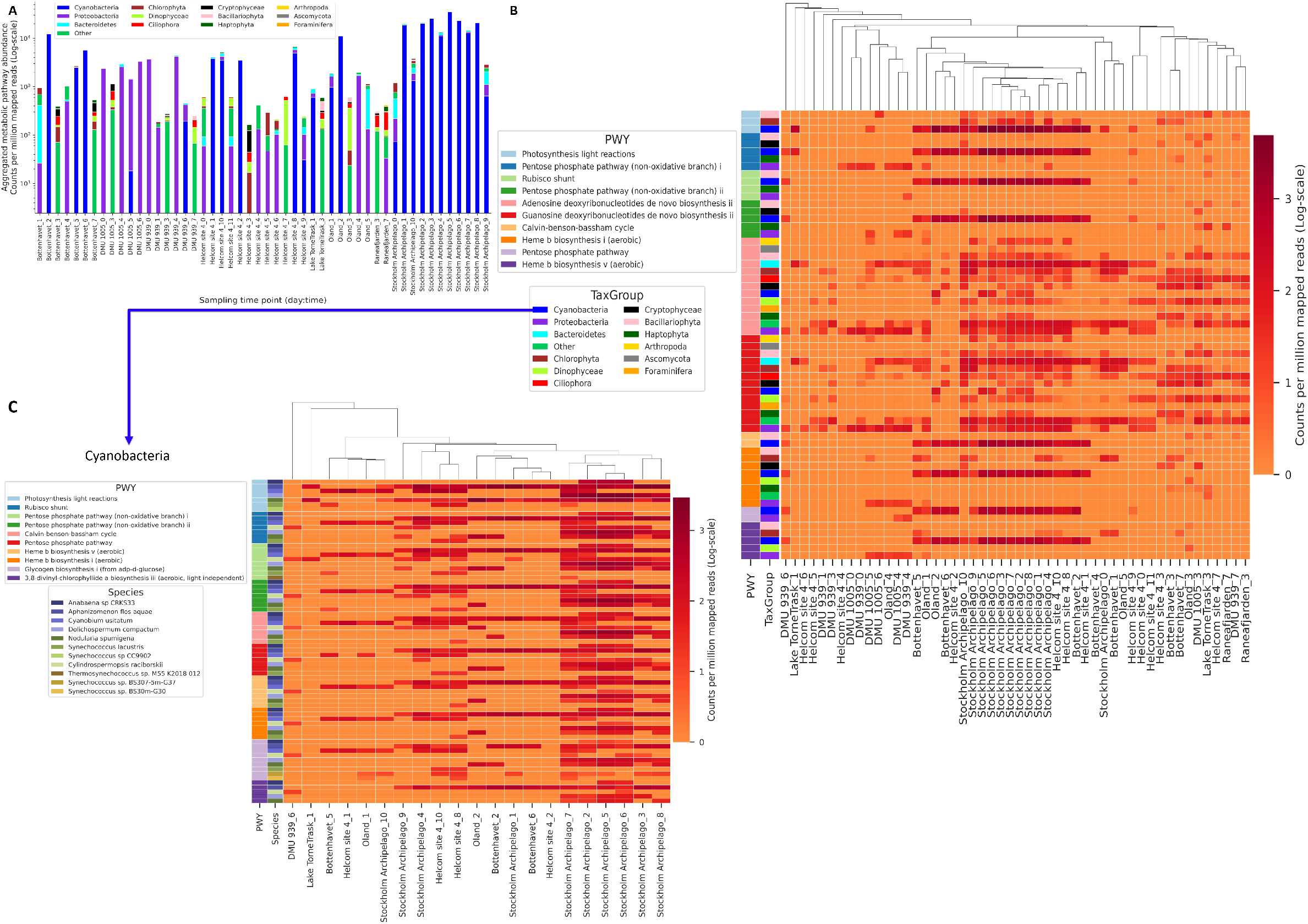
Analysis of metatranscriptome dataset collected at different geographical locations (stations) across the Baltic sea (BioProject record PRJNA320636). A, barcharts illustrating the dominance of major plankton groups in terms of aggregated metabolic activity observed across 178 distinct MetaCyc pathways. B, Heatmap depicting changes (in counts per million mapped reads) across the different stations in the abundance of the top 10 metabolic pathways observed, which were aggregated across the different species belonging to each major taxonomic group analyzed. C, heatmap illustrating changes in the abundance of metabolic pathways observed across stations for each species/strain identified within the Cyanobacteria group only, the dominant taxonomic group observed across most of the samples analyzed.

### Case study 3: Kolody et al. (2019)

In accord with previous findings, our results showed that Cyanobacteria tend to be the globally dominant metabolically active planktonic group in this pelagic ecosystem (Figure 4A), with several other taxonomic groups showing comparatively smaller metabolic activities sparsely through time. Further, we observed that Cyanobacteria tend to show the most consistent periodicity in the activity of the pentose phosphate pathway (alternative pathway to glycolysis for oxidizing glucose and producing NADPH, and C4/C5/C7 sugars), the Rubisco shunt (a highly efficient acetyl-CoA producing pathway), oxygenic photosynthesis and the Calvin-Benson-Bassham cycle (dark reactions) (Figure 4B), which show some degree of disruption in the periodicity at the beginning of the time series recorded, perhaps due to some sort of random alterations in environmental conditions or sampling methodology. Likewise, our results show that energy-producing pathways expressed mostly in Chlorophyta tend to show peak activity levels after midday (2 pm). More generally, when looking at the aggregated metabolic pathway activity profiles across the entire community, using a 2D projection from high dimensional space, we were able to observe a consistent cyclical pattern (Figure 3C). This pattern seems to suggest that global transcriptional profiles of functionally related enzymes tend to be tightly synchronized throughout the day in order to coordinate metabolic responses across the plankton community [57].

**Figure 4.**
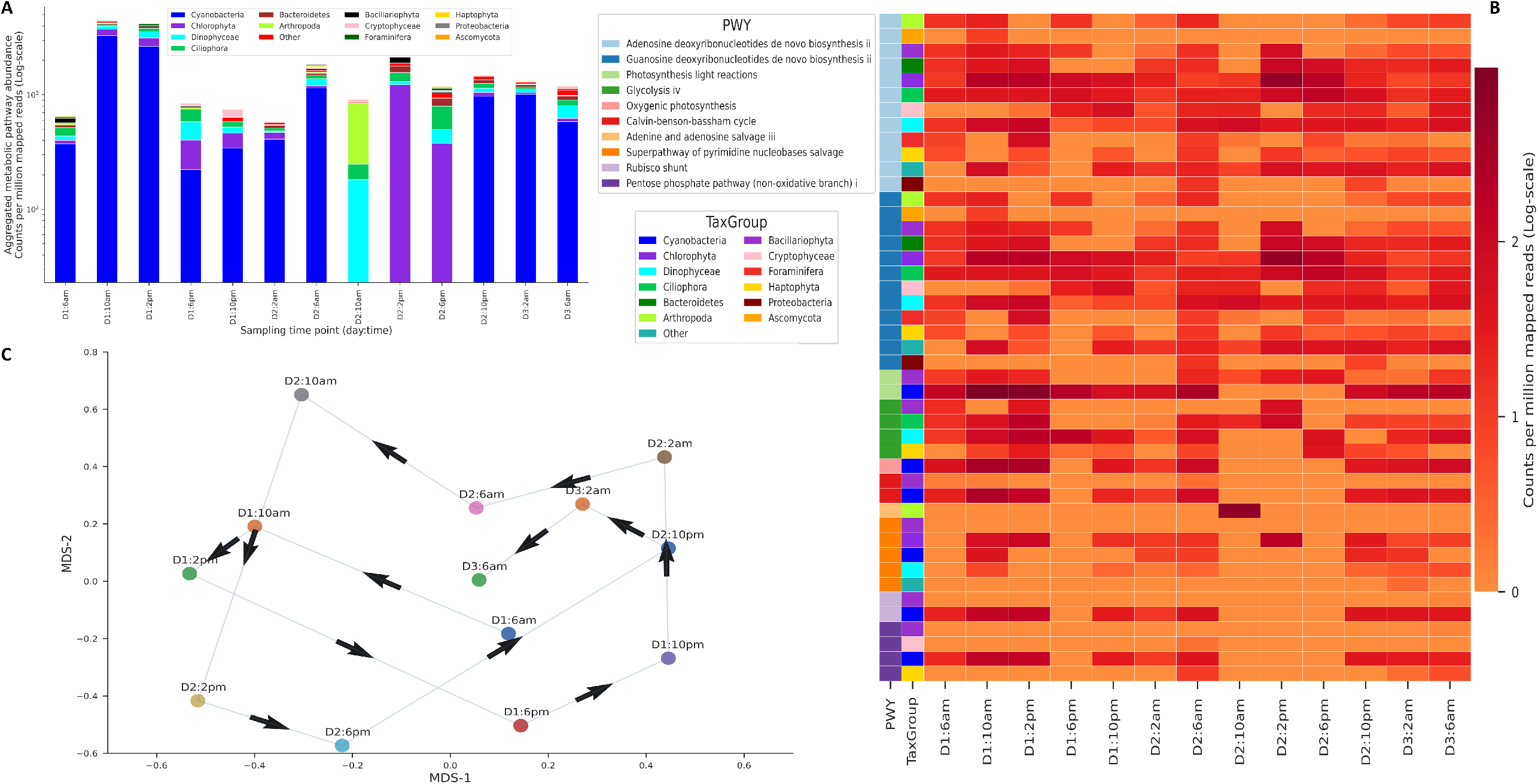
Analysis of time series metatranscriptome dataset collected at different times over nearly 3 consecutive days in surface waters off the Californian coast (BioProject record PRJNA492502). A, barcharts illustrating the dominance of major plankton groups in terms of aggregated metabolic activity observed across 85 distinct MetaCyc pathways. B, Heatmap depicting temporal changes (in counts per million mapped reads) in the abundance of the top 10 metabolic pathways observed, which were aggregated across the different species belonging to each major taxonomic group analyzed. C, projections of the full metabolic pathway abundance profiles using the MDS method applied on the Bray-Curtis pair-wise (species-wise) distance matrix. The arrows in the plot illustrate time directionality of the dataset analyzed.

## DISCUSSION

In this work, we applied the popular microbiome profiling tool HUMANnN on mixed prokaryotic and eukaryotic communities from marine environments. Using a customized plankton reference database we found that the mapping success of transcriptome sequences can be improved by several orders of magnitude. This allowed us to identify and quantify dominant metabolic pathways in largely understudied environments. Congruent with previous observations, we found that database customization can substantially improve our ability to quantitatively study at the systems level, and at various degrees of taxonomic resolution, the physiological status of largely uncharacterized free-living microbiomes [24, 28].

Marine microbial communities are extremely species rich, and full genome sequences of increasingly higher quality are within reach now for major planktonic groups [44]. Unfortunately, our ability to conduct a detailed characterization of the molecular systems mediating species interactions and biogeochemical processes remains so far quite limited, especially for eukaryotic microbial communities. Encouragingly, drastic reductions in sequencing costs are expected [58] and ongoing or planned initiatives for sequencing all of earth’s biodiversity [59] will eventually reduce and later even eliminate this issue.

A particularly challenging problem that emerged from our functional analyses is the necessity to expand the system-based description of microbial networks beyond core metabolic pathways. Future efforts to expand our customized plankton database will be focused on the inclusion of complementary functional annotations, such as the KEGG pathway database [60], as well as databases targeting natural product (secondary metabolite) biosynthetic pathways [61], which will enable a more detailed examination of the relationship between ecological [62] and biogeochemical processes [19, 60]. We will therefore continue to expand on the taxonomic and functional complexity of our customized database with newly released high-quality genome assemblies and gene catalogs and proteomes (e.g. for fungi species). One particularly interesting addition would be the huge protein catalog MERC assembled from metatranscriptome datasets derived from the TARA OCEAN expedition [63], which could be mined for high level representations of the largely unknown sequence space [64] with distant homologies to diverse metabolic functions. This is likely to result in considerable improvements in the mapping success of transcriptome sequences, and thus more comprehensive descriptions of the metabolic pathway activities that drive ecophysiological traits of planktonic ecosystems.

Lastly, a custom database covering a broad taxonomic and functional diversity of a largely unknown environment can facilitate the reverse engineering of molecular system traits suitable for: a) training machine learning models to elucidate statistically significant relationships between core metabolic properties of microbiomes and their surrounding chemical environments; and 2) adequate parameterization of ecosystem biology models with predictive and prescriptive potential on community structure and biogeochemical processes under hypothesized climate change scenarios. In fact, the parameterization of metabolic trait-based models of plankton interaction networks [39, 41] is perhaps the most promising research avenue offered by systems biology-inspired profiling tools such as HUMANnN (3.0). This has the potential to enable a more fine-grained definition of plankton functional types [32, 65] based on aggregated quantities linked to metabolic strategies underlying growth, respiration, inorganic nutrient acquisition, light harvesting, etc. [41]. In this way, self-organization of distinct metabolisms at the community level could be more realistically explored in the context of a global ocean circulation model [66]. This might prove useful to assess whether observed patterns of biogeochemical gradients across space and time can readily emerge from plankton communities with vastly different taxonomic composition and metabolic trait potential [36, 67].

## Acknowledgements

We thank Bart Vanhoorne and Francisco Hernandez for technical support. We also thank Stephane Rombauts and Nicholas Youngblut for their insights and feedback regarding implementation details of some bioinformatic tools used to conduct the study. The resources and services used in this work were provided by the VSC (Flemish Supercomputer Center), funded by the Research Foundation - Flanders (FWO) and the Flemish Government.

## Data Availability

The plankton-specific database reported here is available upon request. Code developed to conduct bioinformatic tasks together with data analyses in the form of Jupyter notebooks are accessible via: https://gitlab.vliz.be/research/sysbioplanktomics.git

## Contributions

JG and PH designed the study. Data gathering and processing were conducted by JG, and the data were analyzed by JG and PH. All authors contributed to the writing of the manuscript.

## Notes

### Competing Interest Statement

The authors have declared no competing interest.

## REFERENCES

1. Cavicchioli R, Ripple WJ, Timmis KN, Azam F, Bakken LR, Baylis M, et al. Scientists’ warning to humanity: microorganisms and climate change. Nat Rev Microbiol 2019; 17: 569–586.

2. Field CB, Behrenfeld MJ, Randerson JT, Falkowski P. Primary production of the biosphere: integrating terrestrial and oceanic components. Science 1998; 281: 237–240.

3. Litchman E, Pinto PT, Edwards KF, Klausmeier CA, Kremer CT, Thomas MK. Global biogeochemical impacts of phytoplankton: a trait□based perspective. Journal of Ecology. 2015: 103: 1384–1396

4. Sokol NW, Slessarev E, Marschmann GL, Nicolas A, Blazewicz SJ, Brodie EL, et al. Life and death in the soil microbiome: how ecological processes influence biogeochemistry. Nat Rev Microbiol 2022.

5. Kour D, Kaur T, Devi R, Yadav A, Singh M, Joshi D, et al. Beneficial microbiomes for bioremediation of diverse contaminated environments for environmental sustainability: present status and future challenges. Environ Sci Pollut Res Int 2021; 28: 24917–24939.

6. Ratzke C, Barrere J, Gore J. Strength of species interactions determines biodiversity and stability in microbial communities. Nature Ecology & Evolution 2020; 4: 376–383.

7. Widder S, Allen RJ, Pfeiffer T, Curtis TP, Wiuf C, Sloan WT, et al. Challenges in microbial ecology: building predictive understanding of community function and dynamics. ISME J 2016; 10: 2557–2568.

8. Lombard F, Boss E, Waite AM, Vogt M, Uitz J, Stemmann L, et al. Globally Consistent Quantitative Observations of Planktonic Ecosystems. Front Mar Sci 2019.

9. Caron DA, Alexander H, Allen AE, Archibald JM, Armbrust EV, Bachy C, et al. Probing the evolution, ecology and physiology of marine protists using transcriptomics. Nat Rev Microbiol 2017; 15: 6–20.

10. Gonzalez E, Pitre FE, Pagé AP, Marleau J, Guidi NW, St-Arnaud M, et al. Trees, fungi and bacteria: tripartite metatranscriptomics of a root microbiome responding to soil contamination. Microbiome 2018; 6.

11. Geisen S, Tveit AT, Clark IM, Richter A, Svenning MM, Bonkowski M, et al. Metatranscriptomic census of active protists in soils. ISME J 2015; 9: 2178–2190.

12. Satinsky BM, Smith CB, Sharma S, Landa M, Medeiros PM, Coles VJ, et al. Expression patterns of elemental cycling genes in the Amazon River Plume. ISME J 2017; 11: 1852–1864.

13. Lambert BS, Groussman RD, Schatz MJ, Coesel SN, Durham BP, Alverson AJ, et al. The dynamic trophic architecture of open-ocean protist communities revealed through machine-guided metatranscriptomics. Proc Natl Acad Sci U S A 2022; 119.

14. Salazar G, Paoli L, Alberti A, Huerta-Cepas J, Ruscheweyh H-J, Cuenca M, et al. Gene Expression Changes and Community Turnover Differentially Shape the Global Ocean Metatranscriptome. Cell 2019; 179: 1068–1083.e21.

15. Plichta DR, Juncker AS, Bertalan M, Rettedal E, Gautier L, Varela E, et al. Transcriptional interactions suggest niche segregation among microorganisms in the human gut. Nat Microbiol 2016; 1: 16152.

16. Abu-Ali GS, Mehta RS, Lloyd-Price J, Mallick H, Branck T, Ivey KL, et al. Metatranscriptome of human faecal microbial communities in a cohort of adult men. Nature microbiology 2018; 3.

17. Borenstein E, Kupiec M, Feldman MW, Ruppin E. Large-scale reconstruction and phylogenetic analysis of metabolic environments. Proc Natl Acad Sci U S A 2008; 105: 14482–14487.

18. Belcour A, Frioux C, Aite M, Bretaudeau A, Hildebrand F, Siegel A. Metage2Metabo, microbiota-scale metabolic complementarity for the identification of key species. Elife 2020; 9.

19. Zhou Z, Tran PQ, Breister AM, Liu Y, Kieft K, Cowley ES, et al. METABOLIC: high-throughput profiling of microbial genomes for functional traits, metabolism, biogeochemistry, and community-scale functional networks. Microbiome 2022; 10: 33.

20. Ibrahim M, Raajaraam L, Raman K. Modelling microbial communities: Harnessing consortia for biotechnological applications. Comput Struct Biotechnol J 2021; 19: 3892–3907.

21. Franzosa EA, McIver LJ, Rahnavard G, Thompson LR, Schirmer M, Weingart G, et al. Species-level functional profiling of metagenomes and metatranscriptomes. Nature Methods 2018; 15: 962–968

22. Beghini F, McIver LJ, Blanco-Míguez A, Dubois L, Asnicar F, Maharjan S, et al. Integrating taxonomic, functional, and strain-level profiling of diverse microbial communities with bioBakery 3. Elife 2021; 10.

23. Caspi R, Billington R, Keseler IM, Kothari A, Krummenacker M, Midford PE, et al. The MetaCyc database of metabolic pathways and enzymes - a 2019 update. Nucleic Acids Res 2020; 48: D445–D453.

24. Youngblut ND, Ley RE. Struo2: efficient metagenome profiling database construction for ever-expanding microbial genome datasets. PeerJ 2021; 9: e12198.

25. Vorobev A, Dupouy M, Carradec Q, Delmont TO, Annamalé A, Wincker P, et al. Transcriptome reconstruction and functional analysis of eukaryotic marine plankton communities via high-throughput metagenomics and metatranscriptomics. Genome Res 2020; 30: 647–659.

26. Sunagawa S, Coelho LP, Chaffron S, Kultima JR, Labadie K, Salazar G, et al. Ocean plankton. Structure and function of the global ocean microbiome. Science 2015; 348: 1261359.

27. Pachiadaki MG, Brown JM, Brown J, Bezuidt O, Berube PM, Biller SJ, et al. Charting the Complexity of the Marine Microbiome through Single-Cell Genomics. Cell 2019; 179: 1623–1635.e11.

28. Karimi E, Geslain E, Belcour A, Frioux C, Aïte M, Siegel A, et al. Robustness analysis of metabolic predictions in algal microbial communities based on different annotation pipelines. PeerJ 2021; 9: e11344.

29. Falkowski PG, Fenchel T, Delong EF. The microbial engines that drive Earth’s biogeochemical cycles. Science 2008; 320: 1034–1039.

30. D’Alelio D, Libralato S, Wyatt T, Ribera d’Alcalà M. Ecological-network models link diversity, structure and function in the plankton food-web. Sci Rep 2016; 6: 21806.

31. Guidi L, Chaffron S, Bittner L, Eveillard D, Larhlimi A, Roux S, et al. Plankton networks driving carbon export in the oligotrophic ocean. Nature 2016; 532: 465–470.

32. Worden AZ, Follows MJ, Giovannoni SJ, Wilken S, Zimmerman AE, Keeling PJ. Environmental science. Rethinking the marine carbon cycle: factoring in the multifarious lifestyles of microbes. Science 2015; 347: 1257594.

33. Villar E, Vannier T, Vernette C, Lescot M, Cuenca M, Alexandre A, et al. The Ocean Gene Atlas: exploring the biogeography of plankton genes online. Nucleic Acids Res 2018; 46: W289–W295.

34. Carradec Q, Pelletier E, Da Silva C, Alberti A, Seeleuthner Y, Blanc-Mathieu R, et al. A global ocean atlas of eukaryotic genes. Nat Commun 2018; 9: 373.

35. Faure E, Ayata S-D, Bittner L. Towards omics-based predictions of planktonic functional composition from environmental data. Nat Commun 2021; 12: 4361.

36. Coles VJ, Stukel MR, Brooks MT, Burd A, Crump BC, Moran MA, et al. Ocean biogeochemistry modeled with emergent trait-based genomics. Science 2017; 358: 1149–1154.

37. Ghyoot C, Flynn KJ, Mitra A, Lancelot C, Gypens N. Modeling Plankton Mixotrophy: A Mechanistic Model Consistent with the Shuter-Type Biochemical Approach. Frontiers in Ecology and Evolution 2017: 5

38. Teodosio MA, Barbosa AMB. Zooplankton Ecology. 2020. CRC Press.

39. Stec KF, Caputi L, Buttigieg PL, D’Alelio D, Ibarbalz FM, Sullivan MB, et al. Modelling plankton ecosystems in the meta-omics era. Are we ready? Mar Genomics 2017; 32: 1–17.

40. D’Alelio D, Eveillard D, Coles VJ, Caputi L, d’Alcalà MR, Iudicone D. Modelling the complexity of plankton communities exploiting omics potential: From present challenges to an integrative pipeline. Current Opinion in Systems Biology 2019: 13: 68–74

41. Mock T, Daines SJ, Geider R, Collins S, Metodiev M, Millar AJ, et al. Bridging the gap between omics and earth system science to better understand how environmental change impacts marine microbes. Glob Chang Biol 2016; 22: 61–75.

42. Horton et al., 2022. World Register of Marine Species. Available from: https://www.marinespecies.orgatVLIZ. DOI: doi:10.14284/170.

43. Delmont TO, Gaia M, Hinsinger DD, Fremont P, Vanni C, Guerra AF, et al. Functional repertoire convergence of distantly related eukaryotic plankton lineages revealed by genome-resolved metagenomics. DOI: https://doi.org/10.1101/2020.10.15.341214.

44. Alexander H, Hu SK, Krinos AI, Pachiadaki M, Tully BJ, Neely CJ, et al. Eukaryotic genomes from a global metagenomic dataset illuminate trophic modes and biogeography of ocean plankton. DOI: https://doi.org/10.1101/2021.07.25.453713.

45. Liu Z, Hu S, Caron D. EukZoo, an aquatic protistan protein database for meta-omics studies. 2018. https://zenodo.org/record/1476236#.YuFBfKhBxPY.

46. Keeling PJ, Burki F, Wilcox HM, Allam B, Allen EE, Amaral-Zettler LA, et al. The Marine Microbial Eukaryote Transcriptome Sequencing Project (MMETSP): illuminating the functional diversity of eukaryotic life in the oceans through transcriptome sequencing. PLoS Biol 2014; 12: e1001889.

47. Hyatt D, Chen G-L, Locascio PF, Land ML, Larimer FW, Hauser LJ. Prodigal: prokaryotic gene recognition and translation initiation site identification. BMC Bioinformatics 2010; 11: 119.

48. Buchfink B, Xie C, Huson DH. Fast and sensitive protein alignment using DIAMOND. Nature Methods. 2015: 12: 59–60.

49. Suzek BE, Wang Y, Huang H, McGarvey PB, Wu CH, UniProt Consortium. UniRef clusters: a comprehensive and scalable alternative for improving sequence similarity searches. Bioinformatics 2015; 31: 926–932.

50. Waterhouse RM, Seppey M, Simão FA, Manni M, Ioannidis P, Klioutchnikov G, et al. BUSCO Applications from Quality Assessments to Gene Prediction and Phylogenomics. Mol Biol Evol 2018; 35: 543–548.

51. Keller O, Kollmar M, Stanke M, Waack S. A novel hybrid gene prediction method employing protein multiple sequence alignments. Bioinformatics 2011; 27: 757–763.

52. Steinegger M, Söding J. Clustering huge protein sequence sets in linear time. Nat Commun 2018; 9: 1–8.

53. BBMap. https://sourceforge.net/projects/bbmap/

54. Cock PJA, Antao T, Chang JT, Chapman BA, Cox CJ, Dalke A, et al. Biopython: freely available Python tools for computational molecular biology and bioinformatics. Bioinformatics 2009; 25: 1422–1423.

55. Elferink S, Wohlrab S, Neuhaus S, Cembella A, Harms L, John U. Comparative Metabarcoding and Metatranscriptomic Analysis of Microeukaryotes Within Coastal Surface Waters of West Greenland and Northwest Iceland. Front Mar Sci 2020.

56. Berg C, Dupont CL, Asplund-Samuelsson J, Celepli NA, Eiler A, Allen AE, et al. Dissection of Microbial Community Functions during a Cyanobacterial Bloom in the Baltic Sea via Metatranscriptomics. Frontiers in Marine Science. 2018: 5.

57. Kolody BC, McCrow JP, Allen LZ, Aylward FO, Fontanez KM, Moustafa A, et al. Diel transcriptional response of a California Current plankton microbiome to light, low iron, and enduring viral infection. ISME J 2019; 13: 2817–2833.

58. Qin Y, Koehler S, Zhao S, Mai R, Liu Z, Lu H, et al. High-throughput, low-cost and rapid DNA sequencing using surface-coating techniques. DOI: https://doi.org/10.1101/2020.12.10.418962.

59. Formenti G, Theissinger K, Fernandes C, Bista I, Bombarely A, Bleidorn C, et al. The era of reference genomes in conservation genomics. Trends Ecol Evol 2022; 37: 197–202.

60. Kanehisa M. KEGG: Kyoto Encyclopedia of Genes and Genomes. Nucleic Acids Research. 2000., 28: 27–30.

61. Navarro-Muñoz JC, Selem-Mojica N, Mullowney MW, Kautsar SA, Tryon JH, Parkinson EI, et al. A computational framework to explore large-scale biosynthetic diversity. Nat Chem Biol 2020; 16: 60–68.

62. Yu JSL, Correia-Melo C, Zorrilla F, Herrera-Dominguez L, Wu MY, Hartl J, et al. Microbial communities form rich extracellular metabolomes that foster metabolic interactions and promote drug tolerance. Nat Microbiol 2022; 7: 542–555.

63. Steinegger M, Mirdita M, Söding J. Protein-level assembly increases protein sequence recovery from metagenomic samples manyfold. Nature Methods 2019; 16: 603–606.

64. Vanni C, Schechter MS, Acinas SG, Barberán A, Buttigieg PL, Casamayor EO, et al. Unifying the known and unknown microbial coding sequence space. Elife 2022; 11.

65. Quere CL, Le Quere C, Harrison SP, Colin Prentice I, Buitenhuis ET, Aumont O, et al. Ecosystem dynamics based on plankton functional types for global ocean biogeochemistry models. Global Change Biology. 2005; 11: 2016–2040.

66. Anderson TR, Follows MJ. Representing Plankton Functional Types in Ocean General Circulation Models: Competition, Tradeoffs and Self-Organizing Architecture. 2010 15th IEEE International Conference on Engineering of Complex Computer Systems. 2010.

67. Follows MJ, Dutkiewicz S, Grant S, Chisholm SW. Emergent Biogeography of Microbial Communities in a Model Ocean. Science. 2007., 315: 1843–1846.

